# Omicron (BA.1) and Sub-Variants (BA.1, BA.2 and BA.3) of SARS-CoV-2 Spike Infectivity and Pathogenicity: A Comparative Sequence and Structural-based Computational Assessment

**DOI:** 10.1101/2022.02.11.480029

**Authors:** Suresh Kumar, Kalimuthu Karuppanan, Gunasekaran Subramaniam

## Abstract

The Omicron variant of the severe acute respiratory syndrome coronavirus 2 (SARS-CoV-2) has now spread throughout the world. We used computational tools to assess the spike infectivity, transmission, and pathogenicity of Omicron (BA.1) and sub-variants (BA.1.1, BA.2, and BA.3) in this study. BA.1 has 39 mutations, BA.1.1 has 40 mutations, BA.2 has 31 mutations, and BA.3 has 34 mutations, with 21 shared mutations between all. We observed 11 common mutations in Omicron’s receptor-binding domain and sub-variants. In pathogenicity analysis, the Y505H, N786K, T95I, N211I, N856K, and V213R mutations in omicron and sub-variants are predicted to be deleterious. Due to the major effect of the mutations characterising, in the receptor-binding domain (RBD), we found that Omicron and sub-variants had a higher positive electrostatic surface potential. This could increase interaction between RBD and electronegative human angiotensin-converting enzyme 2 (hACE2). Omicron and sub-variants had a higher affinity for hACE2 and the potential for increased transmission when compared to the wild type. Among Omicron sub-lineages, BA.2 and BA.3 have a higher transmission potential than BA.1 and BA.1.1. We predicted that mutated residues in BA.1.1 (K478), BA.2 (R400, R490, R495), and BA.3 (R397 and H499) formation of new salt bridges and hydrogen bonds. Omicron and sub-variant mutations at Receptor-binding Motif (RBM) residues such as Q493R, N501Y, Q498, T478K, and Y505H all contribute significantly to binding affinity with human ACE2. Interactions with Omicron variant mutations at residues 493, 496, 498, and 501 seem to restore ACE2 binding effectiveness lost due to other mutations like K417N and Y505H.

## 1 INTRODUCTION

The SARS-CoV-2 RNA genome has been rapidly developing since the first epidemic in late 2019 ^1^. This is mostly due to the virus’s polymerase, which is inherently prone to mistakes, and host immune system selection factors. Many concerns about different mutations in the spike protein surfaced in the previous year. On November 26, 2021, the World Health Organization (WHO) classified Omicron as a variant of concern (VOC) after it was discovered in Botswana. The Omicron variant does have more mutations than prior variants, with many of them occurring in the spike protein’s receptor binding domain ^2^. Despite the spread of several COVID-19 waves over the world, no variant has accumulated mutations or allowed immune evasion to the extent that the SARS-CoV-2 Omicron (B.1.1.529) variant has. The Omicron variant has more than 50 mutations, including 26–32 amino acid substitutions, deletions, and insertions 9 ^3^. The pandemic response strategy for COVID-19 is dependent on the development of treatment options and vaccine formulations ^4–7^. Numerous Omicron mutations, on the other hand, have not been discovered in prior VOCs, and their functional implications have not been thoroughly investigated. However, the discovery of multiple unidentified Omicron mutations inside dominant antibody epitopes has raised worries that vaccination and therapeutic antibody effectiveness may be drastically diminished, necessitating new strategic considerations and research objectives.

A lineage is a set of closely related variations that share a common ancestor, and these may then branch off into sub-lineages, as seems to be occurring with Omicron. The Omicron variant is considered to have divided into four sub-lineages – BA.1, BA.1.1, BA.2, and BA.3 ^8^ which will continue to change in the future. All four lineages were discovered in South Africa at around the same time and location. While all four lineages have expanded worldwide, their rates of spread have varied. This is mostly certainly owing to differences in the spike protein, which is necessary for viral replication and host cell penetration.

Several Omicron lineages have been found after B.1.1.529 was designated as a variation of concern (VOC) on November 26, 2021. Pango lineages BA.1/B.1.1.529.1, BA.1.1/B.1.1.529.1.1, BA.2/B.1.1.529.2 and BA.3/B.1.1.529.3 are among those being monitored by WHO under the ‘Omicron’ umbrella, based on the PANGOLIN (Phylogenetic Assignment of Named Global Outbreak Lineages). According to the outbreak.info website, the cumulative prevalence (the ratio of sequences containing lineage to all sequences submitted to GISAID since lineage identification) for BA.1 is 8 percent (detected in at least 135 countries), BA.1.1 is 5 percent (detected in at least 69 countries), BA.2 is 1 percent (detected in at least 69 countries), and BA.3 is less than 0.5 percent (detected in at least 16 countries) as of 10^th^ February 2022.

It is essential to monitor and study Omicron (BA.1) and its sub-lineages BA.1.1, BA.2, and BA.3, with a particular emphasis on BA.2, for transmissibility, immunological escape qualities, and virulence, which should be prioritised separately and in contrast to BA.1. In our previous study, we compared Omicron (BA.1) and Delta variant of SARS-CoV-2 ^9^. In this study, we compared Omicron variants of BA.1, BA.1.1, BA.2 and BA.3 through computational studies. We studied the sequence and structural characterization of the spike protein which is necessary for viral transmission and entry in these four Omicron variant lineages.

## 2 METHODOLOGY

### 2.1 Omicron and Sub-Variants Protein Sequence retrieval

The FASTA sequence for the Wuhan-HU-1 spike protein was obtained from Uniprot ^10^ (P0DTC2), the protein sequence for BA.2 was obtained from NCBI Genbank ^11^ (UFO69279.1). BA.1.1 (EPI ISL 7605591) and BA.3 (EPI ISL 9092427) genome sequences were retrieved from GSAID ^12^ and translated to protein sequences using the Expasy translate tool. The spike protein sequence for BA.1.1 and BA.3 was selected from the translated protein sequence.

### 2.2 Multiple alignment of Omicron Variants with Wild Type

Using Clustal Omega ^13^ with the default settings, the protein sequence of Wuhan-Hu-1 (Wild type) was aligned with the protein sequences of omicron variant and sub-lineages BA.1, BA.1.1, BA.2, and BA.3. Based on the multiple alignment, mutations were identified. The alignment figure was prepared using Espirt.

### 2.3 Physiochemical characterization

Using the Expasy protparam ^14^, the protein sequences of BA.1, BA.1.1, BA.2, and BA.3’s whole spike protein and RBD were compared to Wuhan-Hu-1 (Wild type). The number of amino acids, molecular weight, theoretical pI, amino acid composition, charged residues, instability index, aliphatic index, and Grand average of hydrophathicity(GRAVY) were all analysed.

### 2.4 Secondary structure and intrinsically disordered prediction

The secondary structure of the Wuhan-Hu-1, Omicron variant (BA.1) and sub-variants (BA.2 and BA.3) was predicted using GOR IV ^15^. The Garnier–Osguthorpe–Robson (GOR) programme analyses secondary protein structure using information theory and Bayesian statistics. The purpose of utilising GOR to combine numerous sequence alignments is to obtain information for enhanced secondary structure discrimination. The Intrinsically disordered of spike protein predicted using PONDR®VLXT tool.

### 2.5 Structural modelling of RBD and Structure analysis

The crystal structure of SARS-CoV-2 spike receptor-binding domain bound with ACE2 of wuhan-HU-1 was obtained using the Protein Data Bank (PDB) (PDB ID: 6M0J) ^16^. The PDB was utilised to get the cryo-electron microscopy (cryo-EM) structure of the SARS-CoV-2 Omicron (BA.1) spike protein in complex with human ACE2 (refinement of RBD and ACE2) (PDB ID:7T9L) ^17^. The structure of BA.1.1, BA.2, and BA.3 was generated by homology modelling with SWISS-MODEL server ^18^ with the BA.1 template (PDB:7T9L). The sequence identity of BA.1.1, BA.2 and BA.3 with template (PDB ID: 7T9l) are 99.50%, 97.01% and 98.51% respectively. Missing residues were inserted to both cyro-EM and homology modelled proteins, and energy was minimised using GROMOS96 43B1 forcefield available at the Swiss-PDB viewer to remove residue steric overlaps at the interface. The Ramachandran plot and Errat plot were used to assess the structure of the homology models. To identify topological and structural changes between wild and Omicron (BA.1, BA.1.1, BA.2 and BA.3) proteins, TM-score and RMSD (root mean square deviation) data were obtained using the I-Tasser online service ^19^. CASTp 3.0 ^20^ was used to predict protein pockets and cavities. The Receptor-binding domain of Omicron and sub-variants were computed for electrostatic potential using electrostatic potential calculated with the Adaptive Poisson–Boltzmann Solver (APBS) program implemented in PyMOL ^21^.

### 2.6 Protein-Protein Docking and stability analysis of RBD-hACE2

The protein-protein docking was carried out between hACE2 and Omicron variants RBD using Hawkdock ^22^ and cluspro ^22^ docking program compared with wild type. For point mutation docking analysis Hex ^23^ docking program was used. We used PDBePISA (PISA) web-based tool to investigate stability of formation of omicron RBD and hACE2 complex.

### 2.7 Pathogenicity analysis

PredictSNP ^24^ was used to determine the pathogenicity of all mutations. Using the PredictSNP web server, prediction algorithms from programmes like as MAPP, PolyPhen1 and PolyPhen-2, SIFT, SNAP, and PANTHER were utilised to achieve a consensus pathogenicity score. The degree of high accuracy is high due to the consensus technique.

## 3 RESULTS AND DISCUSSION

### 3.1 Physio-chemical characterization

Wuhan-Hu-1 (wild type) includes 1273 amino acids, while Omicron BA.1, BA.1.1, BA.2 have 1270 and BA.3 has 1267 amino acids. BA.1, BA.1.1 and BA.2 are three amino acid deficient compared to Wuhan-Hu-1, and BA.3 is six amino acid deficient compared to wildtype. Wuhan-Hu-1 has a molecular weight of 141,178.47, BA.1 has a molecular weight of 141,328.11, BA.1.1 has 141300.09, BA.2 has a molecular weight of 141,185.78, and BA.3 has a molecular weight of 140900.61. BA.3 has a lower molecular weight than Wuhan-Hu-1 owing to the absence of six amino acid residues, but BA.2, BA.1.1 and BA.1 have slightly greater molecular weight than Wuhan-Hu-1 despite the absence of three amino acid residues. A pI number more than 7 indicates that the protein is alkaline, whereas a value less than 7 suggests that the protein is acidic. The theoretical pI of wildtype is 6.24, whereas both BA.1, BA.1.1 have a theoretical pI of 7.14, BA.2 has a theoretical pI of 7.16, and BA.3 has a theoretical pI of 7.35. Omicron BA.1, BA.1.1, BA.2, and BA.3 are shown to be more alkaline than wildtype. According to previous studies, the SARS-CoV-2 S protein is somewhat more positively charged than the SARS-CoV protein, which may result in a greater propensity for binding to negatively charged areas of other molecules through nonspecific and selective interactions ^25^. According to our data, the wild type’s expected charged residues (Arg + Lys) are 103, both BA.1 and BA.1.1 anticipated charged residues are 111, BA.2 predicted charged residues are 108, and BA.3’s predicted charged residues are 109. We noticed that BA.1, BA.1.1, BA.2, and BA.2 all contain significantly more positively charged amino acids (Arg + Lys) than the wild type, which may improve their propensity for binding to negatively charged areas of other molecules such as ACE2. According to the instability index, proteins with a stability score of less than 40 have a stable structure. We observed that all BA.1, BA.1.1, BA.2, and BA.3 (34.21-34.69) are as slight improved stability as compared to wildtype (33.01) ^26^. Aliphatic index, a positive factor associated with improved thermostability of proteins, is 84.67 for wild type, 84.95 for BA.1, BA.1.1, 84.72 for BA.2, and 85.15 for BA.3, indicating that corroborated its stability over a wide range of temperature regime. According to recent study ^27^, increased Omicron thermal stability may result in increased persistence of Omicron in exposed surroundings, posing a greater risk of transmission among household contacts than the Delta form. Increased stability may facilitate viral attachment to host cells by increasing the efficacy of receptor recognition, but it may impair viral membrane fusion.

The grand average of hydropathicity index (GRAVY) was determined using Kyte and Doolittle’s hydropathy values. The hydropathy values range from −2 to +2 for most proteins, with the positively rated proteins being more hydrophobic. GRAVY is projected to be −0.079 for wild type, −0.080 for BA.1 and BA.1.1, −0.074 for BA.2 and −0.071 for BA.3, indicating their hydrophilic nature. BA.1 and BA.1.1 were found to be more hydrophilic, whilst BA.2 and BA.3 were found to be less hydrophilic, the presence of more hydrophobic residues which likely to increase pathogenicity of viral protein ^28^.

From the amino acid composition comparison to the wild type **(Table S1)**, there is an increase in charged residues in the side chains such as Arginine (R) and Lysine (K) in the receptor-binding domain (RBD), which often contributes to the formation of salt bridges ^29^ in BA.1, BA.1.1, BA.2, and BA.3. When compared to the wild type, there is an increase in Asparagine (N) residues in the receptor-binding domain (RBD), which leads to hydrogen bond formation in BA.1, BA.1.1, BA.2, and BA.3. In the receptor binding domains of BA.1, B.A.1.1, BA.2, and BA.3, there is an increase in hydrophobic residues such as phenylalanine (F), Alanine (A), Leucine (L), Proline (P) and leucine (L) compared to the wild type, which are typically buried within the protein core. In BA.1.1, RBD there is increase in polar amino acids like Tryptophan (W) which often at the surface of the protein, Serine (S) which forms hydrogen bond and Valine(V) which are hydrophobic when to compared to all sub-lineages.

### 3.2 Comparison of Omicron Lineages for Unique and Common shared mutations

From the multiple alignment **(Figure S1)** of Omicron and sub-variant with wild type and from four-way Venn-diagram **(Figure 1)** it is inferred that there 39 mutations identified for BA.1, 40 mutations for BA.1.1, 31, mutations for BA.2 and for BA.3 is 34 mutations. Among four-way comparison between Omicron and Sub-variants, for BA.1.1 has one unique mutation R346K, for BA.2 has 8 unique mutation T19I, L24del (deletion), P25del, P26del, A27S, V213G, T376A, R408S and for BA.3 has 1 unique mutation R216del.

**Figure 1.**
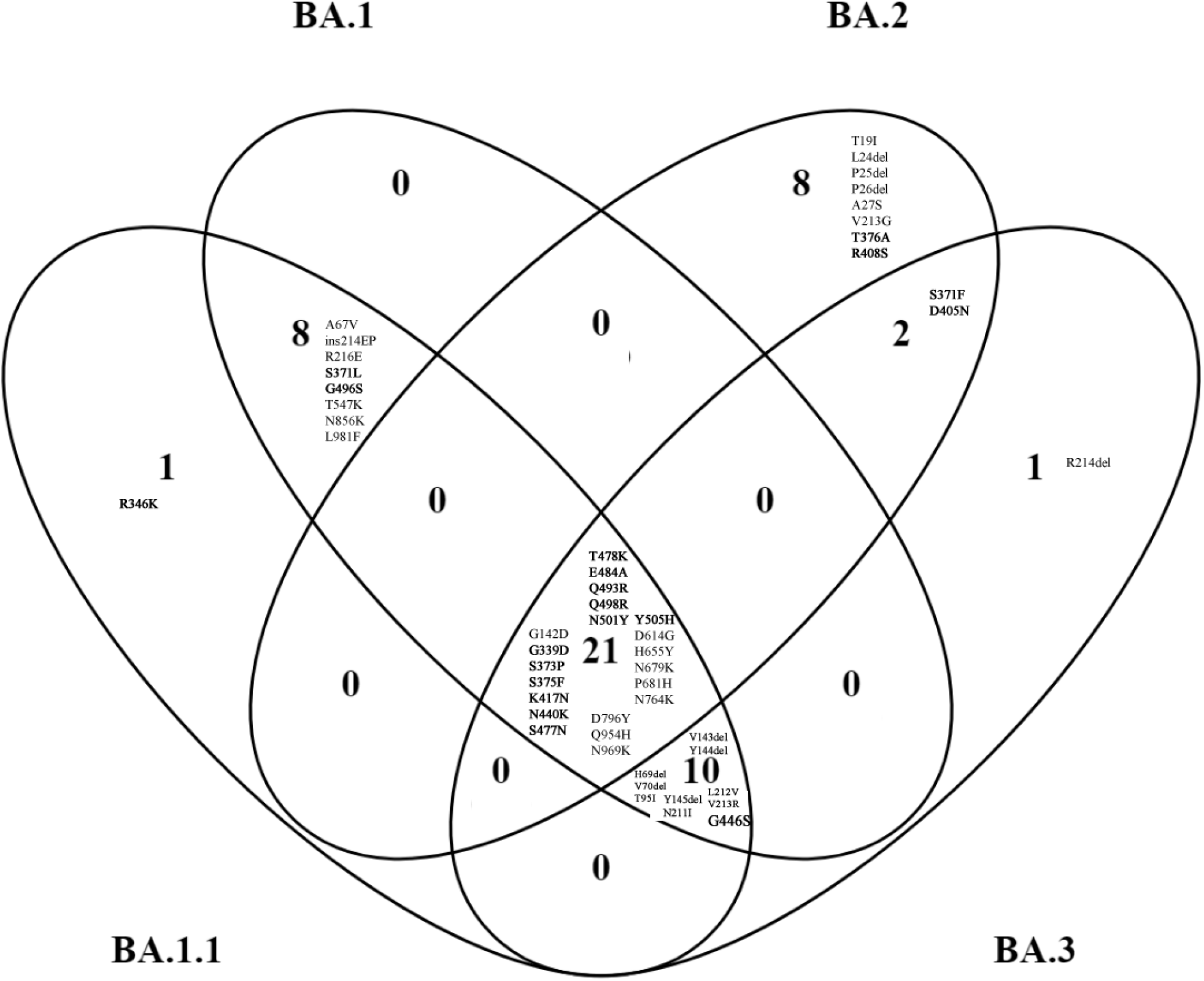
Spike protein mutation in Omicron (BA.1) and Omicron sub-variants (BA1.1, BA.2 and BA.3) compared with four-way Venn-diagram. Receptor-binding domain (residues 319– 541) are marked as bold. Del represents deletion, ins represent insertion.

There are 8 common mutations identified between BA.1.1 and BA.1 are A67V, ins (insertion) 214EP, R216E, S371L, G496S, T547K, N856K, L981F. There are two common identified mutation between BA.2 and BA.3 are S371F and D405N. There are ten common identified mutation between BA.1.1, BA.1 and BA.3 are H69del, V70del, T95I, V143del, Y144del, Y145del. N211I, L212V, V213R, G446S. There are 21 common elements identified between all four Omicron and sub-variants in BA.1.1, BA.1, BA.2 and BA.3 are G142D, G339D, S373P, S375F, K417N, N440K, S477N, T478K, E484A, Q493R, Q498R, N501Y, Y505H, D614G, H655Y, N679K, P681H, N764K, D796Y, Q954H, N969K. Due to triple mutation at the furin cleavage site, such as H655Y, N679K, and P681H Omicron (BA.1) is reported to be extremely transmissible from previous findings ^30^. We observed this triple mutation at the furin cleavage site is common in all sub-variants. There is no commonly shared mutation between BA.1 and BA.2, between BA.1 and BA.3, BA.1, BA.2 and BA.3, between BA.2, BA.3 and BA.1.1, between BA.3 and BA.1.1, between BA.2 and BA.1.1, between BA.1, BA.2 and BA.1.1.

On three-way comparison between BA.1, BA.2 and BA.3, BA.1 has 8 unique mutation A67V, ins214EP, R216E, S371L, G496S, T547K, N856K and L981F. For BA.2, 8 mutation T19I, L24del, P25del, P26del, A27S, V213G, T376A, R408S. BA.3 has one unique mutation R214del. The shared mutation between BA.1 and BA.3 is H69del, V70del, T95I, V143del, Y144del, Y145del, N211I, L212V, V213R, G446S. The shared mutation for BA.2 and BA.3 is S371F, D405N. There are 21 common mutations between BA.1, BA.2 and BA.3 is G142D, G339D, S373P, S375F, K417N, N440K, S477N, T478K, E484A, Q493R, Q498R, N501Y, Y505H, D614G, H655Y, N679K, P681H, N764K, D796Y, Q954H, N969K.

When we compare between BA.1 and BA.2 in two-way comparison, BA.2 has ten unique mutation T19I, L24del, P25del, P26del, A27S, V213G, S371F, T376A, D405N, R408S whereas BA.1 has 18 unique mutation A67V, H69del, V70del, T95I, V143del, Y144del, Y145del, N211I, L212V, V213R, ins214EP, R216E, S371L, G446S, G496S, T547K, N856K, L981F.

In comparison to the RBD (319-541), there are 11 common shared mutation G339D, S373P, S375F, K417N, N440K, S477N, T478K, E484A, Q493R, Q498R, N501Y **(Figure 2A-D)**. Unique mutation for BA.1.1 is R346K, for BA.2 is T376A, R408S. The common shared mutation between BA.1 and BA.1.1 is S371L, G496S.Y505H is the common shared mutation between BA.1, BA.1.1 and BA.2. G446S is the common mutation between BA.1, BA.1.1 and BA.3. S371F, D405N are shared mutation between BA.2 and BA.3. From previous study ^31,32^ it is found that, neutralizing antibodies in infected patients’ sera recognise the receptor-binding domain (319-541 aa) and the N terminal domain of BA.1 and BA.2 (13-306). The divergence suggests that the sub-variants evolved resistance in different immune pressure, possibly in different hosts. All four Omicron lineages has common mutation at Receptor-binding Motif (RBM) region (437-508 a.a) which binds to hACE2 are N440K, S477N, T478K, E484A, Q493R, Q498R, N501Y, Y505H **(Figure 2E)**. There is one unique mutation at RBM for BA.1 at G446S. According to recent study ^33^, during cross-species transmission, SARS-CoV-2 may develop to adapt to a variety of hosts, which inherently favours SARS-CoV-2 evolution. Previously, RBD residues 493, 498, and 501 were identified as key locations for the SARS-CoV-2 host range. Therefore, residues 493, 498, and 501, as well as other changes to omicron’s RBD, are likely to alter the host spectrum of SARS-CoV-2, and the possibility of an omicron variant overcoming the species barrier must be examined further in the future.

**Figure 2.**
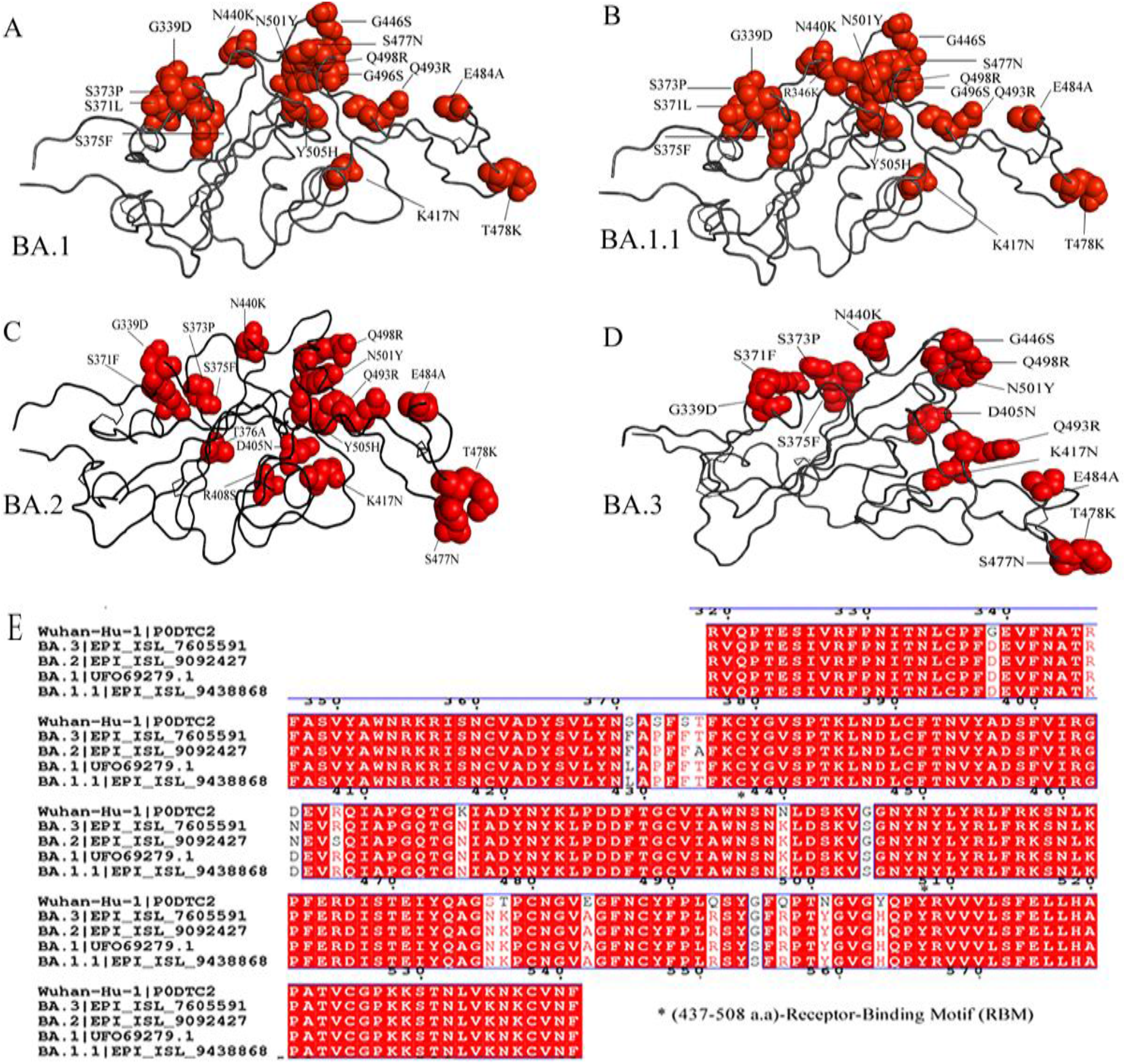
Ribbon diagram of the RBD with residue mutated relative to the wild type.A comparison of **(A)** Omicron-BA.1, Omicron sub-variants **(B)** BA.1.1 **(C)** BA.2 and **(D)** BA.3 mutation in receptor-binding domain (RBD). The multiple alignment **(E)** of RBD showing Receptor-binding motif (RBM) (residues 437-508) of Omicron variants with wild type.

Previous research has shown that deletion of H69/V70 compensates for immune escape mutations that diminish infectivity; thus, it is critical to keep a check out for deletions that have functional consequences ^34^. In contrast to the Omicron BA.1, BA.1.1 and BA.3 spike protein, only the Omicron BA.2 there is absence of the amino acid 69-70 deletion, which is associated with S gene target failure (SGTF) and is not recognised by SGTF ^35^. This complicates tracking its transmission, and some researchers have dubbed it a “stealth” variant. The N501Y mutation, which enhances receptor binding strength, is found in all four BA.1, BA.1.1, BA.2, and BA.3 proteins, and there is a general positive relationship between the stabilising effect of mutations on receptor binding and their population incidence.

### 3.3 Secondary structure and Intrinsically disordered prediction

Through secondary structure prediction **(Table S2)**, when compared to the wild type, Omicron (BA.1, BA.2, and BA.3) entire spike protein and receptor-binding domain exhibit an increase in alpha-helix and a slight decrease in Extended strand. Only the BA.1.1 lineage contains an increased alpha-helix in the entire spike protein, a decreased RBD, and an increased number of extended-strands; as a result, it has only one unique mutation in comparison to BA.1. The projected rise in alpha-helices shows that alpha helices are more prone to mutations than beta strands ^36^. When compared to the wild type, there is a slight decrease in random coil in the total spike protein of all Omicron variants (BA.1) and sub-variants (BA2, BA.3), while there is a slight increase in random coil in the receptor binding domain ^37^.

The Omicron variant of spike protein is less disordered than the wild type, according to the Intrinsically disordered prediction **(Table S3)**. There is a disorder to order transition ^38^ in the receptor-binding motif (468-473) of the Omicron variant, which is required for hACE2 binding. This transition is significant in terms of the effect of disordered residues/regions on spike protein stability and binding to ACE2 ^16^.

### 3.4 Protein structure analysis

For structural comparison, the protein structures of wild type (PDB ID: 6MOJ), Omicron (BA.1) cryo-EM structure (PDB ID: 7T9L), and homology modelled BA.1.1, BA.2 and BA.3 subvariants based on cyro-EM structure (PDB ID: 7T9L) were employed. The structural quality of the homology modelled RBD structures was evaluated using the Ramachandran plot and Errat provided from the SAVES v6.0 webserver (https://saves.mbi.ucla.edu/). Ramachandran’s plot Analyzes residue-by-residue geometry and overall structural geometry to determine the stereochemical quality of a protein structure. Errat compares statistics from highly refined structures to the statistics of non-bonded interactions between distinct atom types and graphs the value of the error function vs location of a 9-residue sliding window. In the Ramachandran plot, the homology modelled of RBD of BA.2 and BA.3 showed residues in most preferred areas with 90.2 percent and residues in extra permitted regions with 9.8 percent. BA.2 and BA.3 both have an Errat score of 89.4. This demonstrates that the overall RBD protein structure of BA.2 and BA.3 is reliable. TM-align tool generates optimum residue-to-residue alignment based on structural similarity using heuristic dynamic programming iterations; scores greater than 0.5 presume the same fold in general. Based on TM-align findings, the RMSD value for all BA.1, BA.1.1, BA.2, and BA.3 is 0.68 based on superposition with wild type. This reveals that Omicron (BA.1, BA.2, and BA.3) have the same structural topology and fold as the wild type. Geometric and topological properties of protein structures, including surface pockets, interior cavities, and cross channels, are of fundamental importance for proteins to carry out their functions. The binding pocket shown in solvent accessible surface area, volume) was predicted for wild type and for Omicron (BA.1, BA.2 and BA.3) using CASTp 3.0. The binding pocket of wild type area is 73.39 and volume is 60.83. For both BA.1, BA.1.1 calculated area is 93.87 and volume is 37.52. For BA.2 predicted area is 49.03 and calculated volume is 16.44. For BA.3, the predicted area is 49.03 and volume predicted is 16.43. From the analysis binding pocket area increased for BA.1 than wild type, while BA.2 and BA.3 decreased whereas binding pocket volume decreases for all BA.1, BA.2 and BA.3 than wild type. We observed that the positive electrostatic surface potential of Omicron and sub-variants is significantly increased as per previous study reported ^39^. This may facilitate RBD interaction with the negatively charged ACE2, hence enhancing the ACE2 receptor’s affinity **(Figure 3)**. Each of these modifications has the potential to affect viral pathogenicity, infectivity, and transmission.

**Figure 3.**
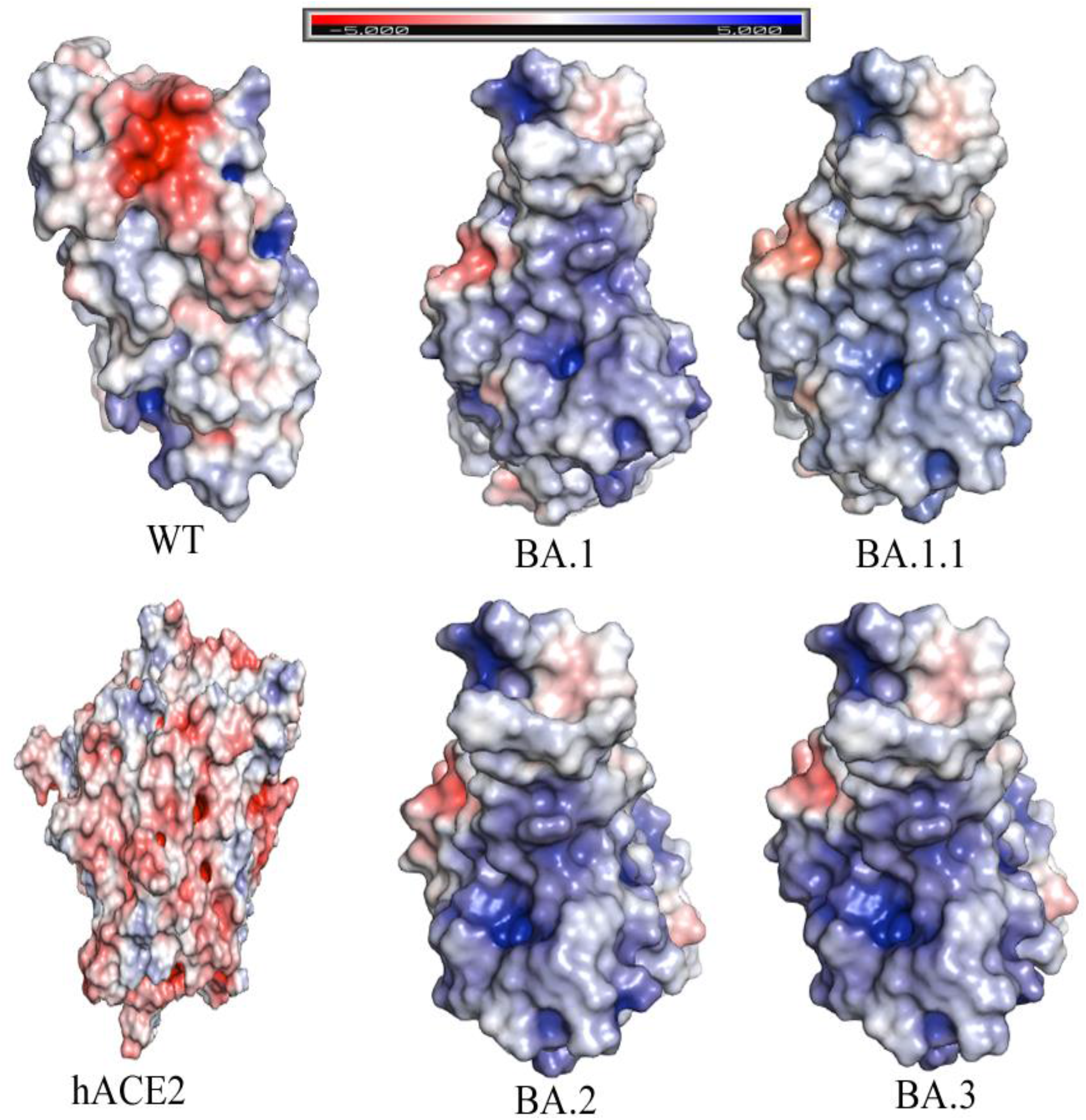
Comparison between the wild-type (WT), Omicron variants (BA.1, BA.1.1, BA.2 and BA.3) spike receptor-binding domains (RBDs) and shown human ACE2. Protein surface is coloured according to the electrostatic potential show in top view. Color scale ranges from −5 kT/e (red) to +5 kT/e (blue) as reported by the bar at the top. The human ACE2 show electronegative potential while Omicron variants shows there is increase of electropositive when compared to wild type.

### 3.5 Pathogenicity and Molecular Docking

The protein sequences were submitted to PredictSNP to predict the effects of mutations on protein function for Omicron SARS-CoV-2 S and hACE2. PredictSNP was used to examine the functional modifications and predictions of the tolerated and deleterious nsSNPs. Our data revealed that among a total of 45 mutations, 39 were neutral, and 6 deleterious with regards to pathogenicity. Y505H, N786K predicted to be deleterious BA.1, BA.1.1, BA.2 and BA.3. T95I, N211I, V213R mutations are predicted to be deleterious in BA.1, BA.1.1, BA.3. N856K mutation are predicted to be deleterious in BA.1, BA.1.1 **(Table 1)**.

**Table 1:** The predicted effect of Spike protein single mutations of Omicron Variants on pathogenicity by using PredictSNP tool. Predicted deleterious mutation marked as bold.

| Wild residue | Position | Target residue | Predicted type | Predict SNP accuracy | Mutation present in |
| --- | --- | --- | --- | --- | --- |
| T | 19 | I | NEUTRAL | 65% | BA.2 |
| A | 27 | S | NEUTRAL | 82% | BA.2 |
| A | 67 | V | NEUTRAL | 73% | BA.1, BA.1.1 |
| **T** | **95** | **I** | **DELETERIOUS** | **60%** | **BA.1, BA.1.1, BA.3** |
| G | 142 | D | NEUTRAL | 74% | BA.1, BA.1.1, BA.3 |
| Y | 144 | D | NEUTRAL | 74% | BA.1, BA.1.1 |
| Y | 145 | D | NEUTRAL | 63% | BA.1, BA.1.1 |
| **N** | **211** | **I** | **DELETERIOUS** | **60%** | **BA.1, BA.1.1, BA.3** |
| L | 212 | V | NEUTRAL | 82% | BA.1, BA.1.1, BA.3 |
| **V** | **213** | **R** | **DELETERIOUS** | **65%** | **BA.1, BA.1.1, BA.3** |
| R | 214 | E | NEUTRAL | 75% | BA.1.1 |
| G | 339 | D | NEUTRAL | 73% | BA.1, BA.1.1, BA.2, BA.3 |
| R | 346 | K | NEUTRAL | 82% | BA.1.1 |
| S | 371 | L | NEUTRAL | 82% | BA.1, BA.1.1, BA.3 |
| S | 371 | F | NEUTRAL | 73% | BA.2 |
| S | 373 | P | NEUTRAL | 82% | BA.1, BA.1.1, BA.2, BA.3 |
| S | 375 | F | NEUTRAL | 75% | BA.1, BA.1.1, BA.2, BA.3 |
| T | 376 | A | NEUTRAL | 65% | BA.2 |
| D | 405 | N | NEUTRAL | 62% | BA.2, BA.3 |
| R | 408 | S | NEUTRAL | 73% | BA.2 |
| K | 417 | N | NEUTRAL | 73% | BA.1, BA.1.1, BA.2, BA.3 |
| N | 440 | K | NEUTRAL | 73% | BA.1, BA.1.1, BA.2, BA.3 |
| G | 446 | S | NEUTRAL | 82% | BA.1, BA.1.1, BA.3 |
| S | 477 | N | NEUTRAL | 82% | BA.1, BA.1.1, BA.2, BA.3 |
| T | 478 | K | NEUTRAL | 63% | BA.1, BA.1.1, BA.2, BA.3 |
| E | 484 | A | NEUTRAL | 82% | BA.1, BA.1.1, BA.2, BA.3 |
| Q | 493 | R | NEUTRAL | 75% | BA.1, BA.1.1, BA.2, BA.3 |
| G | 496 | S | NEUTRAL | 63% | BA.1, BA.1.1 |
| Q | 498 | R | NEUTRAL | 68% | BA.1, BA.1.1, BA.2, BA.3 |
| N | 501 | Y | NEUTRAL | 60% | BA.1, BA.1.1, BA.2, BA.3 |
| **Y** | **505** | **H** | **DELETERIOUS** | **60%** | **BA.1, BA.1.1, BA.2, BA.3** |
| T | 547 | K | NEUTRAL | 82% | BA.1, BA.1.1 |
| D | 614 | G | NEUTRAL | 82% | BA.1, BA.1.1, BA.2, BA.3 |
| H | 655 | Y | NEUTRAL | 73% | BA.1, BA.1.1, BA.2, BA.3 |
| N | 679 | H | NEUTRAL | 82% | BA.1, BA.1.1, BA.2, BA.3 |
| N | 679 | K | NEUTRAL | 82% | BA.3 |
| P | 681 | K | NEUTRAL | 82% | BA.1, BA.1.1, BA.2, BA.3 |
| **N** | **764** | **K** | **DELETERIOUS** | **71%** | **BA.1, BA.1.1, BA.2, BA.3** |
| D | 796 | Y | NEUTRAL | 75% | BA.1, BA.1.1, BA.2, BA.3 |
| **N** | **856** | **K** | **DELETERIOUS** | **75%** | **BA.1, BA.1.1** |
| Q | 954 | H | NEUTRAL | 73% | BA.1, BA.1.1, BA.2, BA.3 |
| N | 969 | K | DELETERIOUS | 54% | BA.1, BA.1.1, BA.2, BA.3 |
| L | 981 | F | NEUTRAL | 82% | BA.1, BA.1.1 |

The interaction with the SARS-CoV-2 virus’s Spike protein receptor binding domain (RBD) and the ACE2 cell surface protein were required for viral infection of cells. The virus increases its evolutionary advantages at the RBD by introducing changes that increase the ACE2-RBD binding affinity or that enable it to avoid antibody detection ^40^. Because the virus’s infectivity in human cells has been improved, any one mutation is unlikely to result in a large increase in viral infectivity. Multiple RBD mutations increase infectivity, which seems to be the case with Omicron, and so appears to be a viable infection pathway. The protein-protein docking method was used, and both HawkDock server and the cluspro docking server was used. The PDB (6M0J) crystal structure of the SARS-CoV-2 spike RBD associated with ACE2 for wild type, as well as the Cryo-EM structure of BA.1 (PDB ID:7T9l) and homology models of BA.2 and BA.3, were used in this investigation.

RBD-ACE2 docking was performed to identify binding pockets and interacting residues using the HawkDock protein-protein docking server. It integrates the ATTRACT docking algorithm, the HawkRank scoring function, and the MM/GBSA free energy decomposition analysis for binding free energy calculations. As shown in **Figure 4**, binding free energy complex for WT is −37.44 (kcal/mol). According to HawkDock server, BA.3 (−73.55) has highest binding free energy compared to BA.2 (−72.36), BA1.1 (−71.56) and BA.1 (−70.6).

**Figure 4.**
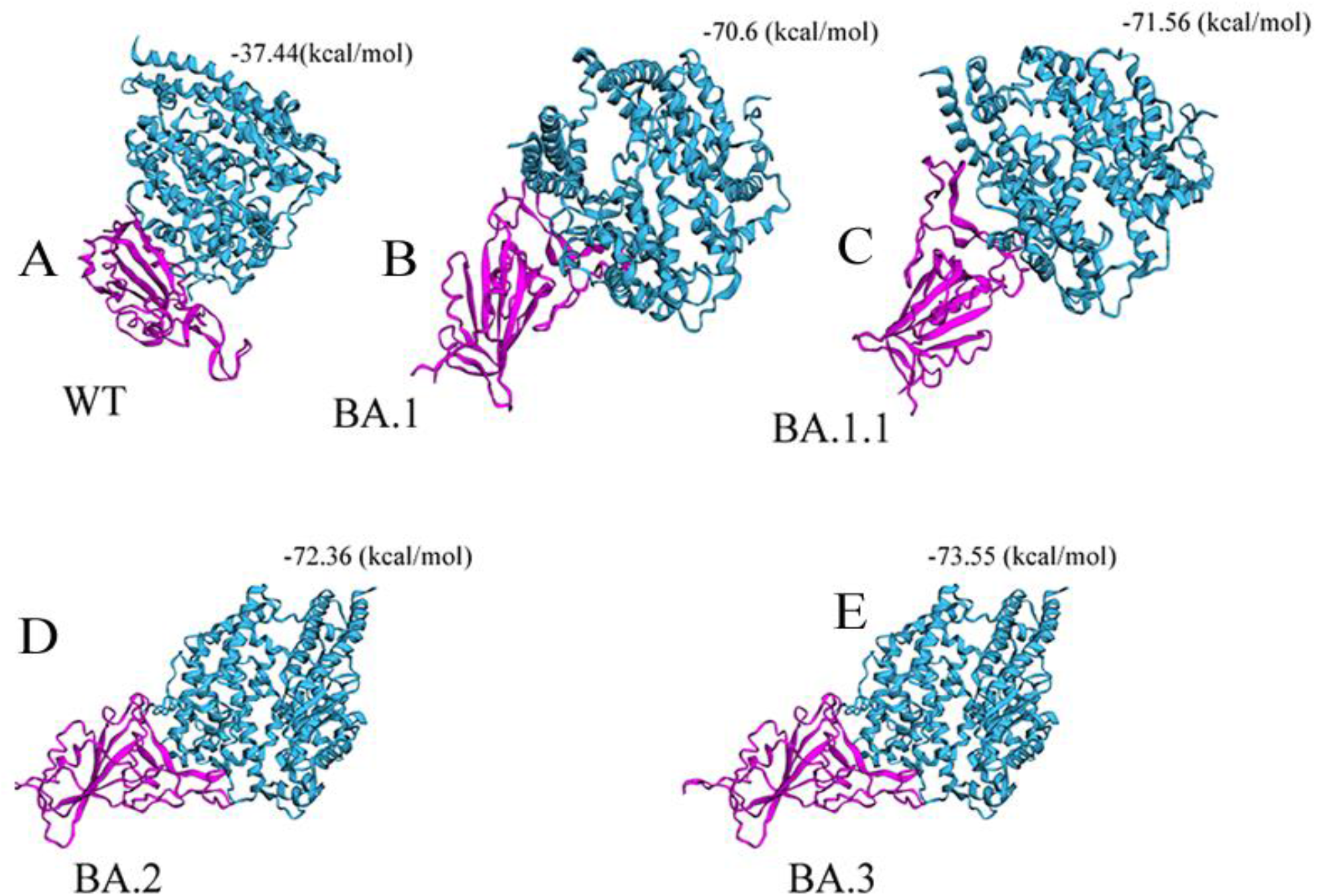
Docking between **(A)** wild-type (WT)-RBD **(B)** Omicron BA.1-RBD, and **(C)** Omicron BA.1.1-RBD **(D)** Omicron BA.2-RBD **(E)** Omicron BA.3-RBD with hACE2. Based on docking energy it shows Omicron BA.2 and BA.3-RBD have a high binding affinity with hACE2 compared to Omicron variant BA.1, B.1.1 and wild-type (WT). Docking score are shown in MM/GBSA free energy decomposition analysis for binding free energy calculations at the top of each variant. hACE2, human angiotensin I-converting enzyme 2; RBD, receptor-binding domain.

The RBD of SARS-CoV-2 was extracted from hACE2 and utilised for protein–protein docking in the wild type and BA.1. After each docking between hACE2 and S protein of each variant, thirty cluster models had been created, of which the model with lowest energy score was selected. The selected cluster models were downloaded in PDB format for further analysis. The homology model is docked with hACE2 for BA.2 and BA.3. The docking energy of WT is −799.6 with hACE2, the docking energy of BA.1 is −943.4 with hACE2, the docking energy for BA.1.1 is −946.8, the docking energy of BA.2 is −974.0 with hACE2, and the docking energy of BA.3 is −999.3 with hACE2. Docking results show that BA.1, BA1.1, BA.2 and BA.3 has a greater affinity for hACE2 than wild type, implying that it has a higher potential for transmission than wild type. We observed among Omicron, BA.2 and BA.3 have a higher affinity for hACE2 than wild type and BA.1, indicating that they had a higher potential for transmission than BA.1 based on both protein-protein docking servers.

In addition, the effect of each point mutation residue of Receptor-binding Motif (RBM) on interaction with hACE2 was studied. Among the common mutations in RBM shared by BA.1, BA.1.1, BA.2, and BA.3, Q493R (−581.53), N501Y (−560.81), Q498 (−527.38), T478K (−517.03), and Y505H had the highest binding affinity with hACE2 (−502.24). Furthermore, the Omicron variant with “Q498R-N501Y” double mutations may enhance RBD binding affinity to the hACE2 receptor. However, the point mutations N440k (−496.38) and E484 (−478.49) had the lowest binding affinity. BA.1 unique mutation G496S (−505.58) has the highest binding affinity. BA.2 unique mutation D405N (−487.86) has the lowest binding affinity with hACE2. Previous research indicated that the N501Y mutation identified in the RBD region increased protein stability as well as a stronger affinity to human ACE2 protein ^41^, which has the potential to increase infectivity and transmission. Despite its location in RBD, S477N exhibits a decreased affinity for the host cell ^42^. Interactions involving Omicron variant mutations at residues 493, 496, 498, and 501 seem to restore ACE2 binding efficiency lost owing to other changes such as K417N and Y505H **(Table 2)**. The retention of total ACE2 binding affinity for the Omicron spike protein shows that compensatory mutations exist that restore increased ACE2 affinity. Omicron has a similar triple mutation like beta variant, “K417N+E484A+N501Y,” which may result in immune escape ^43^.

**Table 2:**
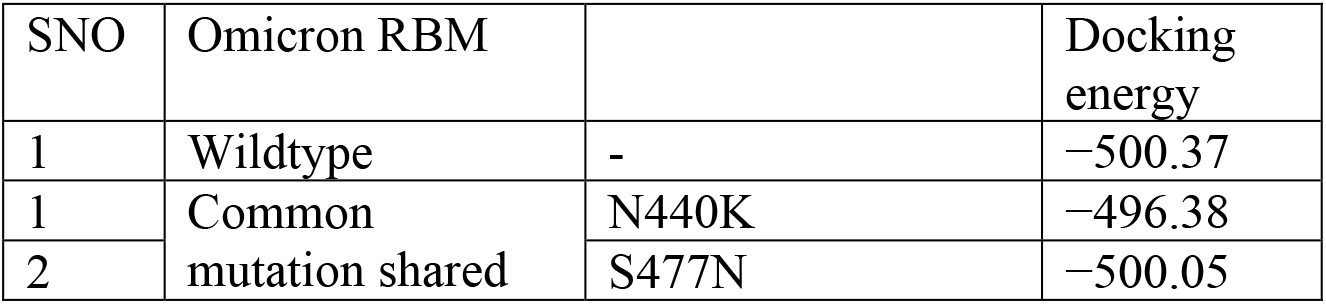
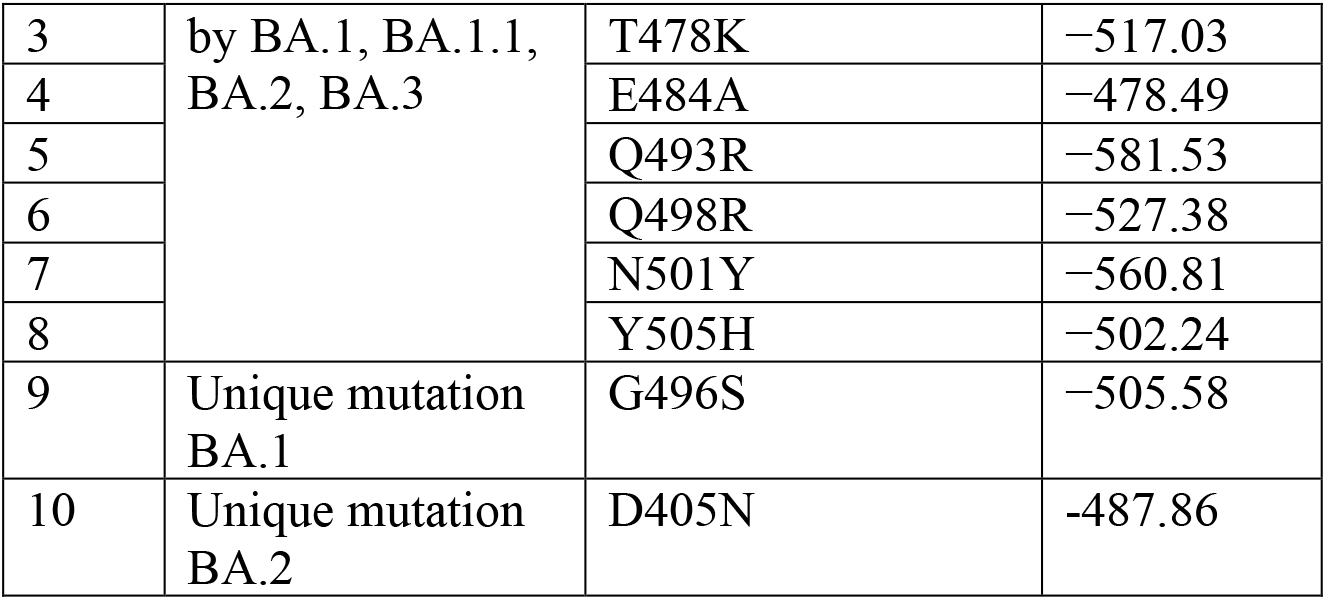
Docking analysis of single-point mutation of Wuhan-RBM (Receptor-binding Motif), Omicron (BA.1)-RBD and Sub-variants (BA1.1., BA.2 and BA.3)-RBD residues with ACE2 using HEX software.

At residue K417, the wild type forms a salt bridge and a hydrogen bond. Cryo-EM structural study of the Omicron variant (BA.1) spike protein in complex with human ACE2 shows new salt bridges and hydrogen bonds produced by R493, R498 mutated residues in the RBD with ACE2. From PISA analysis **(Table 3)**, we predicted that new salt bridges and hydrogen bonds formation for mutated residues in BA.1.1 (K478), BA.2 (R400, R490, R495), and BA.3 (R397 and H499).

**Table 3:** The salt bridge formation and hydrogen bond formation between ACE2-RBD predicted for Omicron and sub-variants using PDBePISA (PISA) web-based tool.

|  | Salt Bridge formation<br>(ACE2-RBD) | Hydrogen Bond formation (ACE2-RBD) |
| --- | --- | --- |
| WT*<br>(PDB ID:<br>6MOJ) | **ASP 30-Lys417** (data from<br>publication) | Gln24-Asn487<br>**Asp30-Lys417**<br>Glu35-Gln493<br>Glu37-Tyr505<br>Asp38-Tyr449<br>Tyr41-Thr500<br>Tyr41-Asn501<br>Gln42-Gly446<br>Gln42-Tyr449<br>Tyr83-Tyr489<br>Tyr83-Asn487<br>Gly502-Lys353<br>Tyr505-Arg393 |
| BA.1*<br>(PDB I<br>D:7T9L) | **GLU 35-ARG 493**<br>**ASP 38-ARG 498** | ASP 38-TYR 449<br>ASP 38-TYR 449<br>GLN 42-TYR 449<br>TYR 83-TYR 489<br>**GLU 35-ARG 493**<br>ASP 38-SER 496<br>ASP 38-ARG 498<br>**GLN 42-ARG 498**<br>TYR 41-THR 500<br>LYS 353-GLY 502<br>HIS 34-TYR 453<br>TYR 83-ASN 487<br>**LYS 353-SER 496**<br>TYR 41-THR 500<br>LYS 353-TYR 501 |
| BA.1.1 | **GLU 57-LYS 478**<br>GLU 35-ARG 490<br>**GLU 35-ARG 493**<br>ASP 38-ARG 490 | LYS 31-TYR 450<br>LYS 68-TYR 470<br>GLN 42-ALA 481<br>GLN 42-CYS 485 |
|  | ASP 30-ARG 495<br>**ASP 30-ARG 498** | THR 27-THR 497<br>ASP 30-TYR 446<br>ASN 61-ASN 474<br>GLU 57-ASN 474<br>GLU 57-ASN 474<br>**GLU 57-LYS 478**<br>GLN 42-ASN 484<br>ASN 61-ASN 484<br>LEU 39-TYR 486<br>**GLU 35-ARG 493**<br>ASP 38-ARG 490<br>**ASP 30-ARG 498** |
| BA.2 | ASP 30-ARG 490<br>**ASP 30-ARG 490**<br>**GLU 35-ARG 495**<br>ASP 38-HIS 502<br>**ASP 38-ARG 400** | ARG 559-ALA 472<br>GLN 388-ASN 484<br>LYS 68-THR 497<br>**ASP 30-ARG 490**<br>**GLU 35-ARG 495**<br>**ASP 38-ARG 400**<br>LYS 353-ASN 414<br>ALA 387-TYR 486 |
| BA.3 | **GLU 35-HIS 499**<br>**GLU 35-ARG 397**<br>GLU 75-ARG 487 | GLN 24-ALA 469<br>LYS 31-TYR 447<br>GLN 42-TYR 495<br>LYS 68-THR 494<br>**HIS 34-ARG 397**<br>ASP 38-VAL 497<br>ASP 38-GLY 498<br>**ASP 38-HIS 499** |

\*Data from publication <sup>16,17</sup>

## 4 CONCLUSION

In this study, Omicron (BA.1) and sub-variants (BA.1.1, BA.2, and BA.3) are compared and investigated with wild type using various computational tools. There are 11 shared common mutations G339D, S373P, S375F, K417N, N440K, S477N, T478K, E484A, Q493R, Q498R and N501Y in RBD Omicron and sub-variants that may contribute significantly to changing the host spectrum of SARS-CoV-2 in immune evasion and potential transmission. The Omicron sub-variants (BA.1.1, BA.2 and BA.3) are likely more transmissible than omicron (BA.1) and Delta. Even if early data indicate that an Omicron infection is less severe than a Delta infection, the quick rise in cases will result in an increase in hospitalizations, placing strain on health care systems to treat individuals with both COVID-19 and other forms of disease. Even though Omicron sub-variants are predicted to be highly transmissible, we anticipate that future COVID-19 waves may be controlled by updated vaccines, reduced vaccine inequity, improved antiviral therapy, and preventative actions taken by the susceptible population.

## 5 LIMITATIONS OF THE STUDY

This evaluation of Omicron variations was mostly confined to computational sequence and structural predictions, and these results should be investigated and validated in future experiments. This study revealed fundamental information about the Omicron protein structures, laying the framework for future research on the SARS-CoV-2 Omicron and sub-variants.

## ACKNOWLEDGMENTS

The authors acknowledge Faculty of Health and Life Sciences, Management and Science University, Malaysia; Division of Cardiovascular Medicine, Radcliffe Department of Medicine, Wellcome Centre for Human Genetics, University of Oxford, Oxford, UK and Department of Physiology, Anatomy, and Genetics, University of Oxford, Oxford, UK. The authors also acknowledge GISAID (https://www.gisaid.org/) for facilitating open data sharing.

## AUTHOR CONTRIBUTIONS

Conceptualization, investigation, writing—original draft preparation: Suresh Kumar. Writing—review and editing: Kalimuthu Karuppanan and Gunasekaran Subramaniam. Data curation and formal analysis: Suresh Kumar. Methodology, data curation, investigation, and validation: Suresh Kumar, Kalimuthu Karuppanan, Gunasekaran Subramaniam. Project administration, and supervision: Suresh Kumar.

