## Supplementary Information for "Omicron (BA.1) and Sub-Variants (BA.1, BA.2 and BA.3) of SARS-CoV-2 Spike Infectivity and Pathogenicity: A Comparative Sequence and Structural-based Computational Assessment"

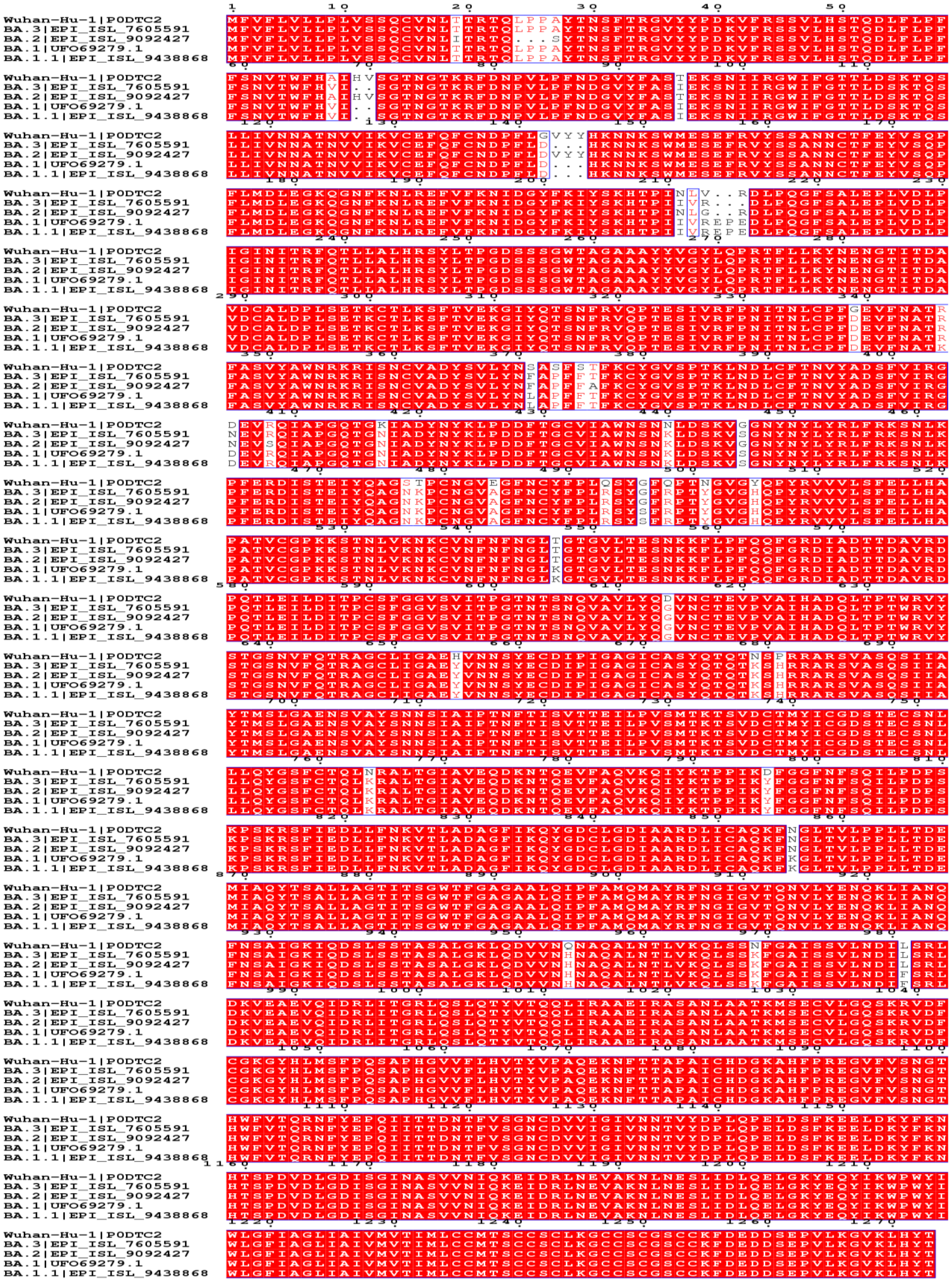

**Figure S1** Multiple alignment of Omicron variants with Wuhan-Hu-1 (Wild type)

**Table S1**: Amino acid composition comparison between Omicron BA.1, BA.1.1, BA.2 and BA.3 with reference to wild type (wuhan-Hu-1)

| | **Wuhan-Hu-1 -Whole Spike** | **Omicron-Whole Spike (BA.1)** | **Omicron-Whole Spike (BA.1.1)** | **Omicron-Whole Spike (BA.2)** | **Omicron-Whole Spike (BA.3)** | **Wuhan-Hu-1 RBD** | **Omicron-RBD (BA.1)** | **Omicron-RBD (BA.1.1)** | **Omicron-RBD (BA.2)** | **Omicron-RBD (BA.3)** |
|---|---|---|---|---|---|---|---|---|---|---|
| **Ala (A)%** | 6.2 | 6.2 | 6.2% | 6.3% | 6.2% | 5.2% | **5.7%** | **6.1%** | **6.1%** | **5.7%** |
| **Arg (R)%** | 3.3 | 3.5 | 3.4% | 3.4% | 3.5% | **4.8** | **5.7%** | **6.5%** | **5.2%** | **5.7%** |
| **Asn (N)%** | 6.9 | 6.5 | 6.5% | 6.8% | 6.7% | **9.2%** | **10.0%** | **10.7%** | **10.5%** | **10.5%** |
| **Asp (D)%** | 4.9 | 4.9 | 4.9% | 4.8% | 4.8% | 3.9% | 4.4% | 2.8% | 3.9% | 3.9% |
| **Cys (C)%** | 3.1 | 3.1 | 3.1% | 3.1% | 3.2% | 3.9% | 3.9% | 4.2% | 3.9% | 3.9% |
| **Gln (Q)%** | 4.9 | 4.6 | 4.6% | 4.6% | 4.7% | 3.1% | 2.2% | 1.9% | 2.2% | 2.2% |
| **Glu (E)%** | 3.8 | 3.9 | 3.9% | 3.7% | 3.7% | 3.1% | 2.6% | 3.3% | 2.6% | 2.6% |
| **Gly (G)%** | 6.4 | 6.2 | 6.2% | 6.5% | 6.3% | 6.6% | 5.7% | **4.2%** | **6.6%** | **6.1%** |
| **His (H)%** | 1.3 | 1.4 | 1.4% | 1.5% | 1.4% | 3.1% | 0.9% | 0.9% | 0.9% | 0.9% |
| **Ile (I)%** | 6.0 | 6.1 | 6.1% | 6.1% | 6.2% | 3.9% | 3.9% | 4.2% | 3.9% | 3.9% |
| **Leu (L)%** | 8.5 | 8.4 | 8.4% | 8.4% | 8.4% | **6.1%** | **7.0%** | **6.1%** | **6.6%** | **6.6%** |
| **Lys (K)%** | 4.8 | 5.3 | 5.4% | 5.1% | 5.1% | 5.2% | **6.1%** | **7.0%** | **5.7%** | **5.7%** |
| **Met (M)%** | 1.1 | **1.1** | **1.1%** | **1.1%** | **1.1%** | 0.0% | 0.0% | 0.0% | 0.0% | 0.0% |
| **Phe (F)%** | 6.0 | 6.2 | 6.2% | 6.2% | 6.2% | **7.0%** | **7.9%** | **7.9%** | **8.3%** | **8.3%** |
| **Pro (P)%** | 4.6 | 4.6 | 4.6% | 4.4% | 4.6% | **5.7%** | **6.1%** | **6.5%** | **6.1%** | **6.1%** |
| **Ser (S)%** | 7.8 | 7.6 | 7.6% | 7.6% | 7.6% | 7.4% | 6.6% | 7.0% | 6.1% | 6.1% |
| **Thr (T)%** | 7.6 | 7.4 | 7.4% | 7.4% | 7.5% | 5.7% | 5.2% | 5.1% | 5.2% | 5.7% |
| **Trp (W)%** | 0.9 | 0.9 | 0.9% | 0.9% | 0.9% | 0.9% | 0.9% | 1.4% | 0.9% | 0.9% |
| **Tyr (Y)%** | 4.2 | 4.3 | 4.3% | 4.4% | 4.3% | 6.6% | 6.6% | 4.7% | 6.6% | 6.6% |
| **Val (V)%** | 7.6 | 7.6 | 7.6% | 7.6% | 7.6% | 8.7% | 8.7% | 9.3% | 8.7% | 8.7% |
| **Pyl (O)%** | 0.0 | 0.0% | 0.0% | 0.0% | 0.0% | 0.0% | 0.0% | 0.0% | 0.0% | 0.0% |
| **Sec (U)%** | 0.0 | 0.0% | 0.0% | 0.0% | 0.0% | 0.0% | 0.0% | 0.0% | 0.0% | 0.0% |

**Table S2:** Secondary structure prediction and comparison of Omicron (BA.1, B.A.1.1, BA.2, BA.3) variant and sub-variants with reference to wild type (Wuhan-Hu-1)

| | **Wuhan-Hu-1-whole Spike** | **Omicron-Spike (BA.1)** | **Omicron Spike (BA.1.1)** | **Omicron Whole Spike (BA.2)** | **Omicron Whole Spike (BA.3)** | **Wuhan -Hu-1- RDB** | **Omicron-RDB (BA.1)** | **Omicron-RDB (BA.1.1)** | **Omicron-RBD (BA.2)** | **Omicron-RBD (BA.3)** |
|---|---|---|---|---|---|---|---|---|---|---|
| **Alpha helix (Hh)** | 21.52% | 23.46% | 22.91% | 23.31% | 23.68% | 6.55% | 8.30% | 5.61% | 8.73% | 8.30% |
| **Extended strand (Ee)** | 22.07% | 20.55% | 20.87% | 20.87% | 20.92% | 22.71% | 18.34% | 23.36% | 18.34% | 20.09% |
| **Random coil (Cc)** | 56.40% | 55.98% | 56.22% | 55.83% | 55.41% | 70.74% | 73.36% | 71.03% | 72.93% | 71.62% |

**Table S3**: Intrinsically disordered prediction using PONDR®VLXT tool.

| | No.of residues disordered | Overall percent disordered | Predicted disorder segment | Number Disordered Regions |
|---|---|---|---|---|
| Wuhan-HU-1 | 98 | 7.70 | [17]-[20]<br>[468]-[475]<br>[601]-[608]<br>[672]-[709]<br>[869]-[871]<br>[945]-[950]<br>[982]-[986]<br>[992]-[994]<br>[1023]-[1023]<br>[1174]-[1194]<br>[1264]-[1264] | 11 |
| Wuhan-RBD | 18 | 7.86 | [317]-[322]<br>[468]-[473] | 2 |
| Omicron (BA.1) | 85 | 6.69 | [17]-[20]<br>[208]-[221]<br>[598]-[607]<br>[675]-[706]<br>[867]-[868]<br>[1020]-[1020]<br>[1171]-[1191]<br>[1261]-[1261] | 8 |
| Omicron-RBD (BA.1) | 6 | 2.62 | [317]-[322] | 1 |
| Omicron (BA.1.1) | 85 | 6.69 | | 8 |
| Omicron RBD (BA1.1) | 6 | 2.62 | [317]-[322] | 1 |
| Omicron (BA.2) | 78 | 6.14 | [406]-[407]<br>[598]-[607]<br>[675]-[706]<br>[866]-[868]<br>[979]-[983]<br>[989]-[991]<br>[1020]-[1020]<br>[1171]-[1191]<br>[1261]-[1261] | 9 |
| Omicron RBD (BA.2) | 8 | 3.49 | [317]-[322]<br>[408]-[409] | 2 |
| Omicron (BA.3) | 80 | 6.31 | [17]-[20]<br>[595]-[604]<br>[672]-[703] | 9 |

| | | | | |
|---|---|---|---|---|
| | | | [863]-[865]<br>[976]-[980]<br>[986]-[988]<br>[1017]-[1017]<br>[1168]-[1188]<br>[1258]-[1258] | |
| Omicron RBD (BA.3) | 6 | 2.62 | [317]-[322] | 1 |
